# Dynamic production and loss of flagellar filaments during the bacterial life cycle

**DOI:** 10.1101/767319

**Authors:** Xiang-Yu Zhuang, Shihao Guo, Zhuoran Li, Ziyi Zhao, Seiji Kojima, Michio Homma, Pengyuan Wang, Chien-Jung Lo, Fan Bai

## Abstract

Bacterial flagella are large extracellular protein organelles that drive bacteria motility and taxis in response to environmental changes. Previous research has focused mostly on describing the flagellar assembly, its rotation speed and power output. However, whether flagella are permanent cell structures and, if not, the circumstances and timing of their production and loss during the bacterial life cycle remain poorly understood. Here we used the single polar flagellum of *Vibrio alginolyticus* as our model and, using *in vivo* fluorescence imaging, revealed that the percentage of flagellated bacteria (PFB) in a population varies substantially across different bacterial growth phases. In the early-exponential phase, the PFB increases rapidly in respect to incubation time, mostly through widespread flagella production. In the mid-exponential phase, the PFB peaks at around 76% and the partitioning of flagella between the daughter cells is 1:1 and strictly at the old poles. After entering the stationary phase, the PFB starts to decline, mainly because daughter cells stop making new flagella after cell division. Interestingly, we discovered that bacteria can actively abandon flagella after prolonged stationary culturing, though cell division has long been suspended. Lack of glucose was found to be a major factor promoting flagellar disassembly. We also revealed that the active loss of flagella was initiated by breakage in the rod connecting the extracellular filament to the basal body formed by MS- and C-rings. Our results highlight the dynamic production and loss of flagellar filaments during the bacterial life cycle.

## Main Text

The flagellum of a flagellated bacterium is a remarkable structure consisting of a reversible rotary motor, a short proximal hook, and a thin helical filament [1]. Since the flagellar system was discovered, this delicate nanomachine has attracted much research and, while some aspects of the flagellar system have been well studied, others have not [2-6]. Meanwhile, flagella are not only critical for bacterial motility, their constituent flagellins are also important antigens that can stimulate both the innate inflammatory response and the development of adaptive immunity [7-9]. Two specialized receptors on immune cells, cell surface Toll-like receptor 5 (TLR5) [10-12] and intracellular receptor Ipaf [13, 14], are responsible for recognizing flagellins as a warning of a pathogenic bacterial invasion. Knowing when and how bacteria produce and lose flagella, especially under stressed conditions, is valuable for understanding the way bacteria either trigger or evade host immunity surveillance, a central topic in pathogen-host interactions.

Previous research has shown that bacterial flagella are hollow protein cylinders with 20 nm outer and 2 nm inner diameters [15]. A flagellum is assembled from the inside out, beginning with a basal body embedded in the cell membrane and completing with the structural components of the rod, hook, and filament sequentially unfolded and exported by a type III secretion system [1, 16, 17]. These subunits pass through the flagellum’s nascent central channel and crystallize at its tip, extending the growing flagellum, which ultimately reaches about 3-10 times the length of the cell body [15]. Harnessing the transmembrane electrochemical proton motive force (PMF), each bacterial flagellar motor (BFM) [18, 19] quickly rotates its flagellum, subsequently propelling the bacterial cell body at a speed of 15-100 µm/s [20]. Using cutting-edge biophysical methods, the torque-speed relationship [21-25], stepping [26], and switching [27, 28] of the BFM have been investigated. Very recently, the dynamic assembly of flagellar filaments has been characterized in real time [29-31]. However, questions still remain about whether flagella are permanent structures of the cell and, if not, the timing of their production and loss during different phases of the bacterial life cycle remain poorly understood.

As for the flagellar basal structure, it is composed of the ring structures, which are C-ring, MS-ring, and LP-ring, and the rod which penetrates and connects the rings [32-34]. MS-ring, which is composed of ca. 30 subunits of membrane protein FliF, is assembled at first in the flagellar formation [35]. C-ring, which is composed of three kind proteins, FliG, FliM and FliN, is assembled beneath MS-ring to interact N-terminal domain of FliG and the C-terminal domain of FliF [36, 37]. The flagellar outer membrane complexes, which seems to contain LP-ring, and many incomplete flagellar subassemblies have been observed by cryo-electron tomography in various bacteria [32, 38, 39]. The polar flagellum of *Vibrio alginolyticus* has extra ring structures around LP-ring, called T-ring and H-ring [33, 40].

Using the single polar flagellum of *V. alginolyticus* as the model, we systematically surveyed the dynamic production and loss of flagella across different growth phases of the bacterial life cycle. Implementing fast fluorescent labeling and imaging, we were able to track the location and timing of flagellar growth with high spatial and temporal resolutions. We showed that flagella are not permanent cellular structures. Instead, the probability that a cell might possess a flagellum varies substantially across different growth phases. Interestingly, we revealed that bacteria can actively abandon flagella after prolonged stationary culturing, and depletion of their glucose was found to be the factor triggering this disassembly of flagella. Our study illustrates a dynamic allotting of resources and energy for flagellar construction during the bacterial life cycle and describes a new flagellar disassembly process model.

## Results

### Percentage of flagellated *V. alginolyticus* varies in different growth phases

In our experiment, an overnight culture of *V. alginolyticus* was diluted 1:100 and continued to grow in VC growth medium. Because *V. alginolyticus* flagella are each covered by sheath which is believed to be an extension of the bacterial cell membrane, we can readily label the flagella with a lipophilic fluorescent dye (FM 4-64) and visualize whether a given cell has a polar flagellum (Figure 1A) [31].

**Figure 1.**
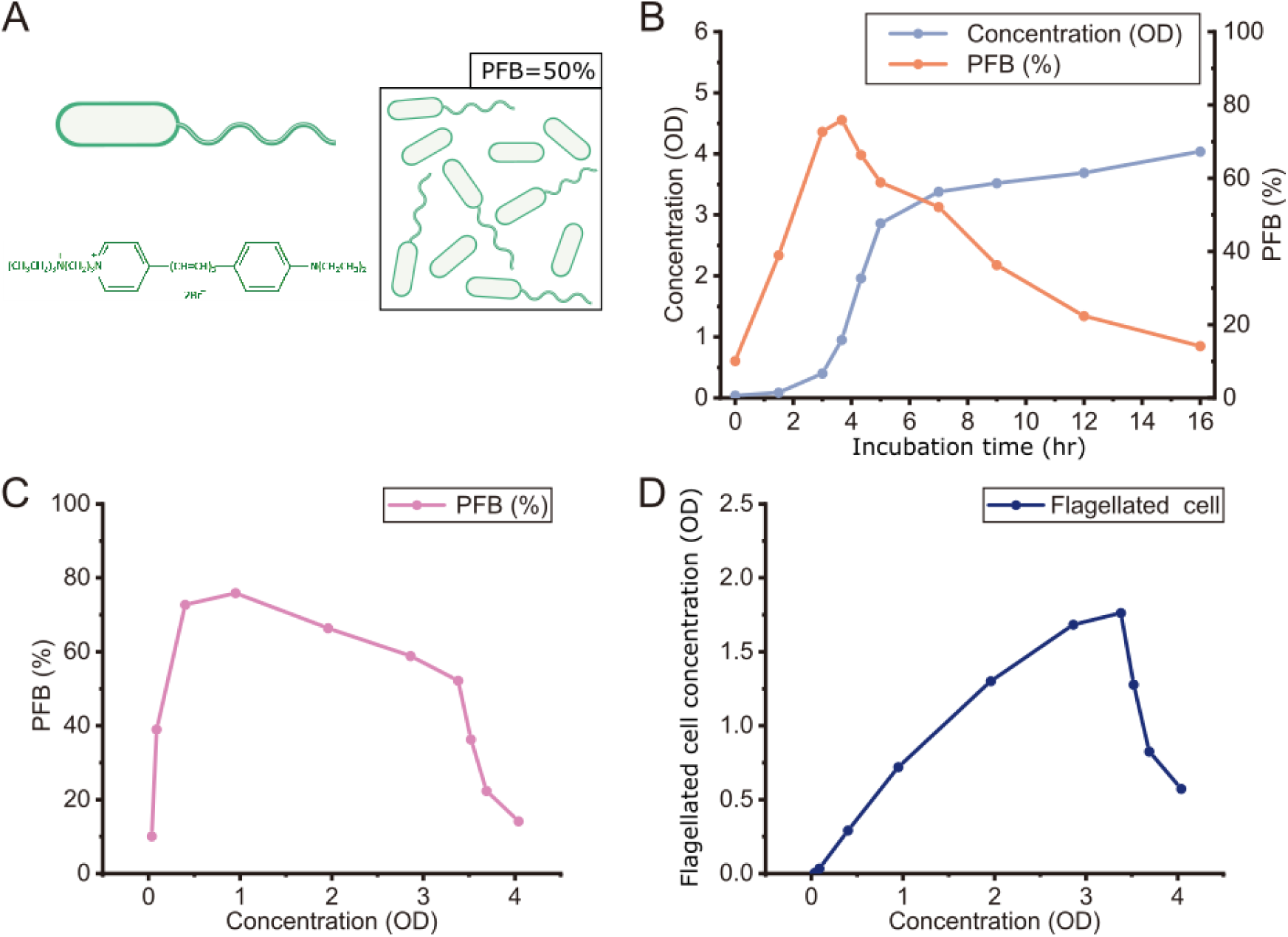
Dynamic measurement of the percentage of flagellated bacteria (PFB) of *V. alginolyticus* during different growth phases. (A) A schematic illustration of the fluorescent labeling of *V. alginolyticus*’s polar flagella with lipophilic fluorescent dye FM 4-64 (Materials and Methods) and how the PFB was measured. (B) The dynamic relationship between the PFB and incubation time suggested that bacterial flagella are not permanent structures of the cell. At least 300 cells were counted for each time point. (C) The relationship between the PFB and cell concentration implied that flagellar production and loss followed 4 different periods: rapid production, stable construction, dilution, and active loss. (D) The sharp decline in the relationship between absolute flagellated cell number and cell concentration indicated that flagella are widely lost in the stationary phase. OD, optical density.

After growing in VC medium, a 100 μL aliquot of bacterial culture was taken from the flask at an indicated time point and examined under a fluorescence microscope. This procedure continued for the other indicated time points. For each, we scanned the imaging field and determined the percentage of flagellated bacteria (PFB), which was defined as the number of cells with fluorescent flagella divided by the total number of cells examined (Figure 1A and Materials and Methods).

At the population level, we saw that the PFB differed greatly across different phases of bacterial growth (Figure 1B). After overnight culturing, most cells lost their flagella, perhaps a result of nutrient depletion in the medium, resulting in a PFB of only about 10%. But, when we reinoculated the bacteria into fresh VC growth medium and the subsequent bacterial growth was in its early-exponential phase (0-3 hrs of incubation), the PFB increased dramatically (Figure 1B), peaking at 76% in the midexponential phase (3-3.67 hrs of incubation). As we continued to culture the bacteria, the PFB started to decline after reaching the late-exponential phase (5 hrs of incubation). Once the cells began the stationary phase, the PFB dropped sharply and finally returned to about 10% (Figure 1B). We confirmed the flagella lost from cell in the late-exponential phase to detect the filament proteins (Fig. S1).

To further understand the dynamics of flagellar production and loss, we plotted the PFB versus cell concentration (Figure 1C) and calculated the flagellated cell concentration (cell OD × PFB) with respect to the total cell concentration (cell OD), as shown in Figure 1D. Our findings indicated that in the early-exponential phase, many non-flagellated cells produced new flagella, resulting in a marked increase in both the PFB and flagellated cell number. In the mid-exponential phase, the PFB peaked while flagellated cell numbers kept increasing. In the late-exponential phase, the PFB declined while flagellated cell numbers remained constant, indicating cell division without creation of new flagella. When the culture entered its stationary phase, the PFB dropped further and the cells actively abandoned flagella, which was seen in the sharp decline in flagellated cell number (Figure 1C, D).

### In the early-exponential phase flagella are widely produced

Following our finding that the PFB increased rapidly in the early-exponential phase of bacterial growth, we examined the mechanism underlying these dynamic PFB changes at the single cell level.

To investigate the flagellar growth pattern in this phase, we again used FM 4-64 to stain the flagella and conducted time-lapse recording under a fluorescent microscope. Overnight, wild-type *V. alginolyticus* strain VIO5 was transferred into fresh VC medium (see Materials and Methods) at the ratio of 1:100 for regrowth. After 0-3 hours, bacteria cells were observed under a fluorescent microscope and flagellar growth was recorded in real-time for 180 minutes.

We also constructed a VIO5 derivative strain (NMB344) that expressed enhanced green fluorescent protein-fused FliG (EGFP-FliG) (see Materials and Methods) [41, 42]. FliG is a component protein in the part of the C-ring structure which is responsible for torque generation with stator units on the basal body of the bacterial flagellar motor, where the C-ring noncovalently attaches to the MS-ring. According to previous studies, the C-ring must form before the flagellum can assemble [1, 43]. Therefore, using this strain we can validate the production of a new flagellum by identifying a localized EGFP-FliG spot at the base of the flagellum.

Initial microscopic observations showed that most bacteria possessed no flagella, a result consistent with the population measurement. Very soon though, we saw a widespread production of flagellar stubs and the PFB increased rapidly. During this process, many bacterial cells without flagella became flagellated, accompanied by formation of localized EGFP-FliG spots at cell poles. The appearance of flagella mostly followed 2 patterns: (I) A flagellum was made at one pole of the bacterium before cell elongation and division (Figure 2A, I). (II) Following cell division, a flagellum was made in one daughter cell (Figure 2A, II). Both mechanisms could account for the increase in the PFB and the flagellated cell number that we observed in the population measurements (Figure 1C, D). Representative time-lapse images showing the production of new flagella labelled by FM 4-64 and emergence of EGFP-FliG spots (not on the same cell) are shown in Figure 2B, C.

**Figure 2.**
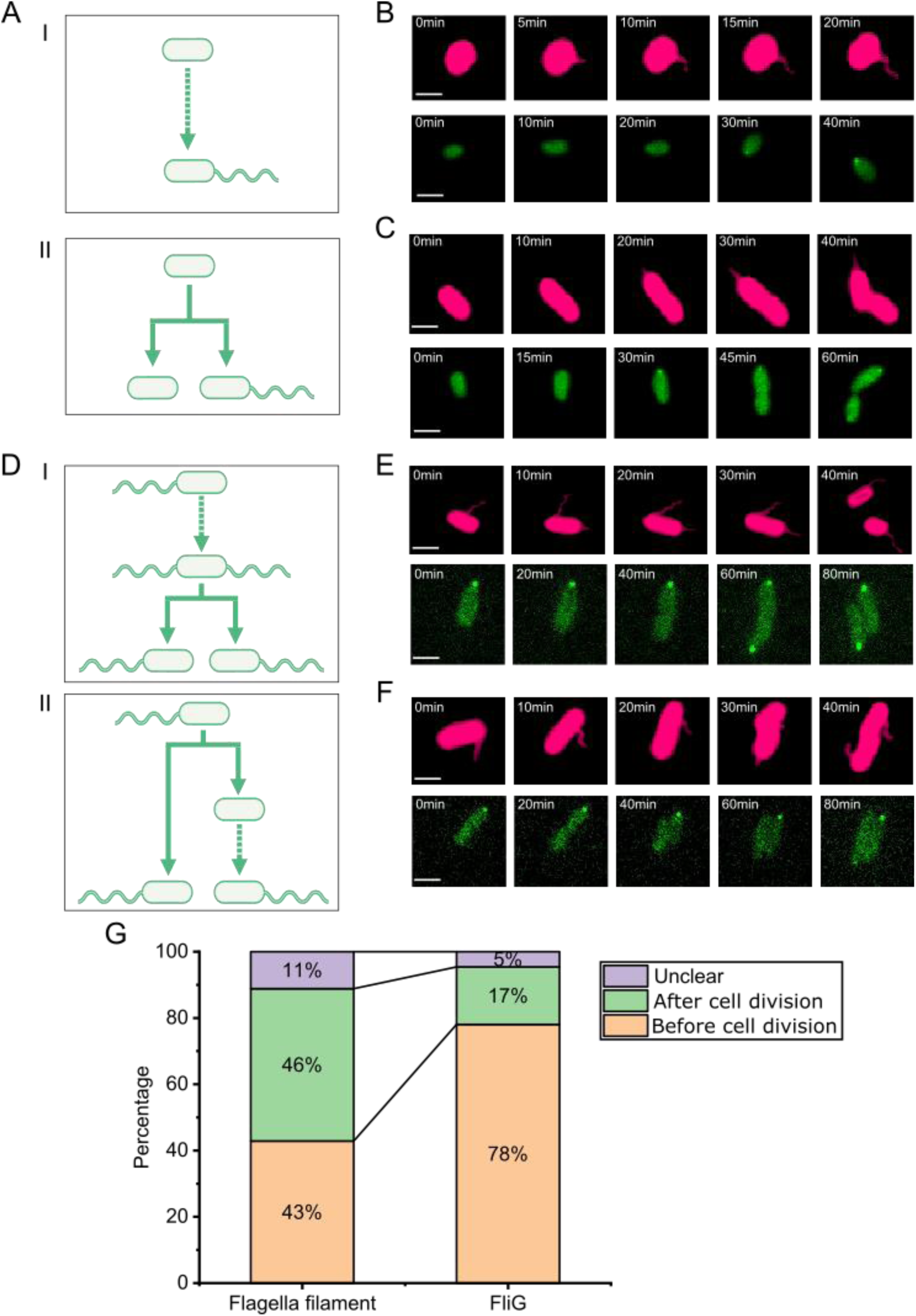
Time-lapse fluorescent imaging revealed the temporal and spatial patterns of flagella production in *V. alginolyticus* during the early-(A) and mid-exponential (D) phases of its growth. (A)(D) Cartoon schematics showing different scenarios for flagellar production. (A) I. Flagellum was made directly from one pole of the bacterium before cell elongation and division. II. Following cell division, 1 daughter cell produced a new flagellum. (D) I. A second flagellum is produced before cell division. II. A new flagellum is produced after cell division. (B)(C) In the early-exponential phase, flagella production was prolific. (E)(F) In the mid-exponential phase, new flagella were made at 1:1 ratio and only at the old poles during cell division. Red images showing new flagella are of FM 4-64 dyed cells and green images of strain NMB344 cells contain EGFP-FliG spots. (G) Percentages of flagella filament production and FliG cluster formation before cell division vs. after cell division.

### In the mid-exponential phase new flagella are made at 1:1 ratio and strictly at the old poles during cell division

In the mid-exponential phase (3.67 hrs), the PFB peaked at about 80%. During this stage, bacteria grew and divided vigorously, and most cells possessed flagella. Many flagellated bacteria generated daughter cells, all with polar flagella after cell division, so the PFB stayed constant at about 80% (Figure 1B). Then we studied the localization and timing of newly grown flagella among different cells.

During bacterial cell division, each daughter cell has an old pole and a new pole. The old poles existed in the mother cell prior to cell division, while the new poles form only after cell division is complete. Through real-time single-cell imaging, we found that new flagella were uniformly made at the old poles during cell division (Figure 2D). The appearance of flagella mostly followed 2 patterns: (I) A new flagellum was made at the pole opposite that of the pre-existing flagellum, then cell division soon followed (Figure 2D, I). (II) Cell division first generated a daughter cell without a flagellum, but a new flagellum soon formed at the cell’s old pole (Figure 2D, II). These mechanisms maintained the high PFB and increased flagellated cell numbers. Representative time-lapse images showing the production of new flagella labelled by FM4-64 and the emergence of EGFP-FliG spots at flagellar bases (not on the same cell) are shown in Figure 2E, F.

In our investigation of the temporal sequence between the production of new flagella and cell division, we defined cell division completion as the time point when the septum splits revealing 2 separate cells. The time when flagella growth initiates was defined as the time point when a new flagellum was identified. By staining both flagella and cell body, we found that new flagella could grow either before (43%) or after (46%) cell division was finished (Figure 2G). However, the temporal sequence between EGFP-FliG cluster assembly and cell division differed in that FliG cluster assembly could occur either before (78%) or after (17%) cell division was complete (Figure 2G). This observed higher percentage of FliG cluster formation before cell division agrees with the fact that FliG cluster formation is a prerequisite of flagellar growth.

### In the late-exponential phase, daughter cells from cell division stop to make new flagella

When bacterial growth entered the late-exponential phase (5 hrs), cell growth and division slowed and the PFB began to decline (Figure 1B). In this phase, bacteria were still dividing but the daughter cells made no new flagella, resulting in a ‘diluted’ PFB. This decline in the PFB followed 2 patterns: (I) Most flagellated bacteria divided into 2 daughter cells, with one retaining the flagellum and the other never making a new flagellum (Figure 3A, I). (II) The bacteria without flagella continued to divide and the resulting daughter cells had no flagella (Figure 3A, II). Both mechanisms served to decrease the PFB, but the flagellated cell number remained constant. Representative time-lapse images showing the cell division of both bacteria with and without pre-existing flagella (labelled by FM4-64) are shown in Figure 3B, C.

**Figure 3.**
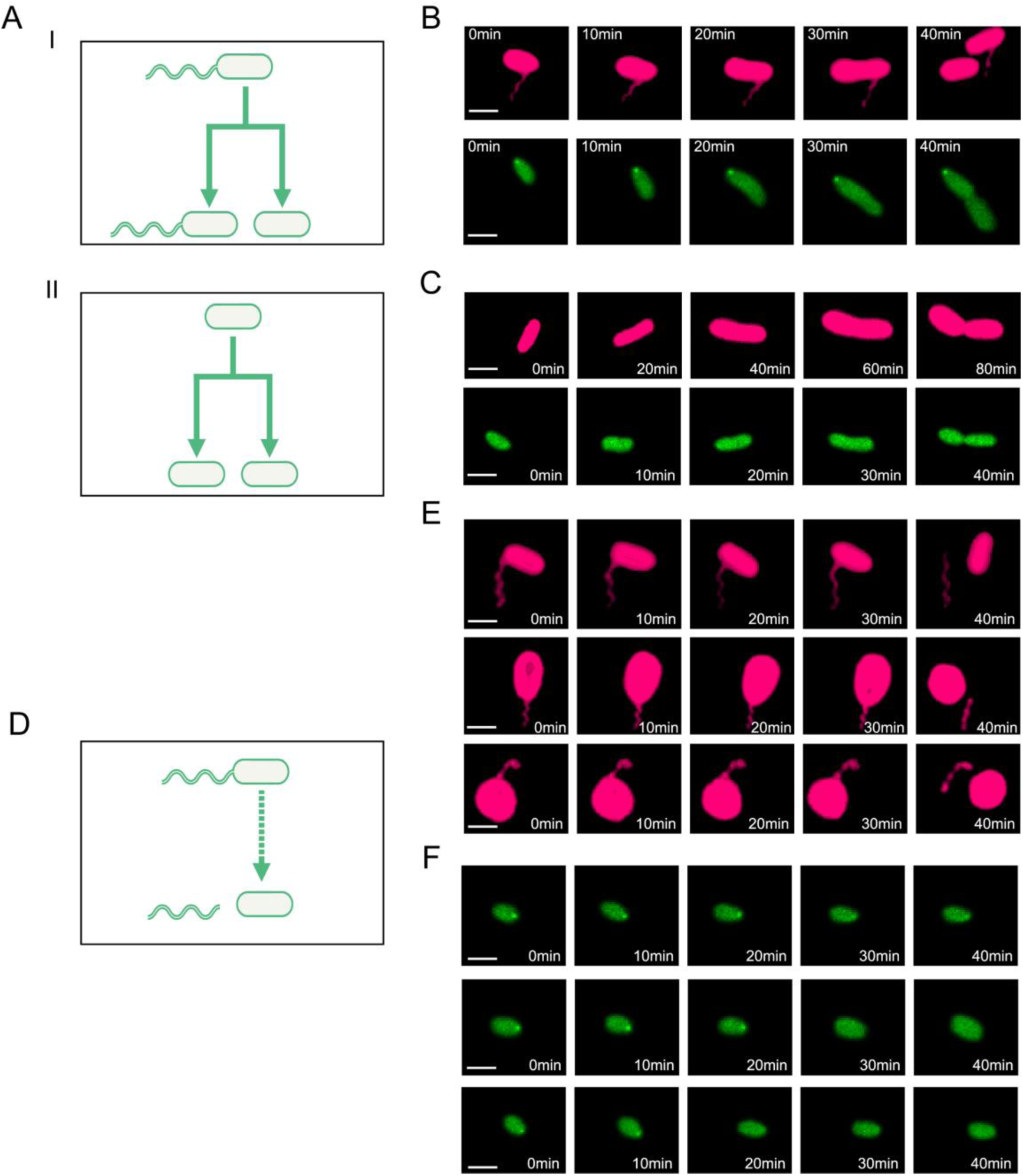
Time-lapse fluorescent imaging revealed the spatial and temporal patterns of flagellar loss in *V. alginolyticus* during the late-exponential (A) and stationary (D) cell growth phases. (A)(D) Cartoon schematics showing different scenarios for flagellar loss. (A) I. Following fission, 1 cell retains the flagella and the other fails to grow one. II. Cells without flagella divide and both daughters fail to produce flagella. (D) Cells with flagella shed their flagella. (B)(C) In the late-exponential phase, flagella production ceases during cell division. Red images showing flagella are of FM 4-64-dyed cells and green images of strain NMB344 cells contain EGFP-FliG spots. (E)(F) In the stationary phase, cells actively abandoned flagella.

### In the stationary phase, *V. alginolyticus* actively abandon flagellar filaments

When bacterial growth entered the stationary phase (7+ hrs), the bacteria’s OD was basically stable, indicating that bacterial cell numbers did not increase. However, the PFB continued decreasing during this phase (Figure 1B), suggesting that cell division was not the only mechanism through which bacteria lose flagella.

Under the microscope, we observed the bacteria that had been transferred to fresh VC medium at 10 hrs after overnight culture and found that bacteria that shed their flagella frequently also showed no signs of growth (Figure 3D). Representative time-lapse images of cells labeled with FM4-64 showing flagella loss and the disappearance of EGFP-FliG spots (not on the same cell) are shown in Figure 3E, F. These phenomena explained the continued PFB and flagellated cell number declines after the bacteria entered the stationary phase, in which cell division had ceased.

### Lack of glucose promotes flagellar disassembly

We next sought to understand what factors triggered *V. alginolyticus* flagellar loss in its stationary phase. In the stationary phase, depleted nutrients in the growth medium and accumulated metabolites may have contributed to flagellar loss. To test the hypothesis that nutrient depletion precipitated active flagella abandonment, we used glucose-less TMN medium supplemented with glucose of varying concentrations to treat bacteria and flagellated cells were counted under a microscope. As shown in Figure 4A, the PFB in a medium with no glucose decreased rapidly to 5% in 5 hrs. However, the addition of 5 mM glucose slowed down the PFB drop to 30% in 5 hrs and 15 mM glucose was enough to stop the loss of flagella. Therefore, our results suggest that glucose is important for bacterial flagella retention.

**Figure 4.**
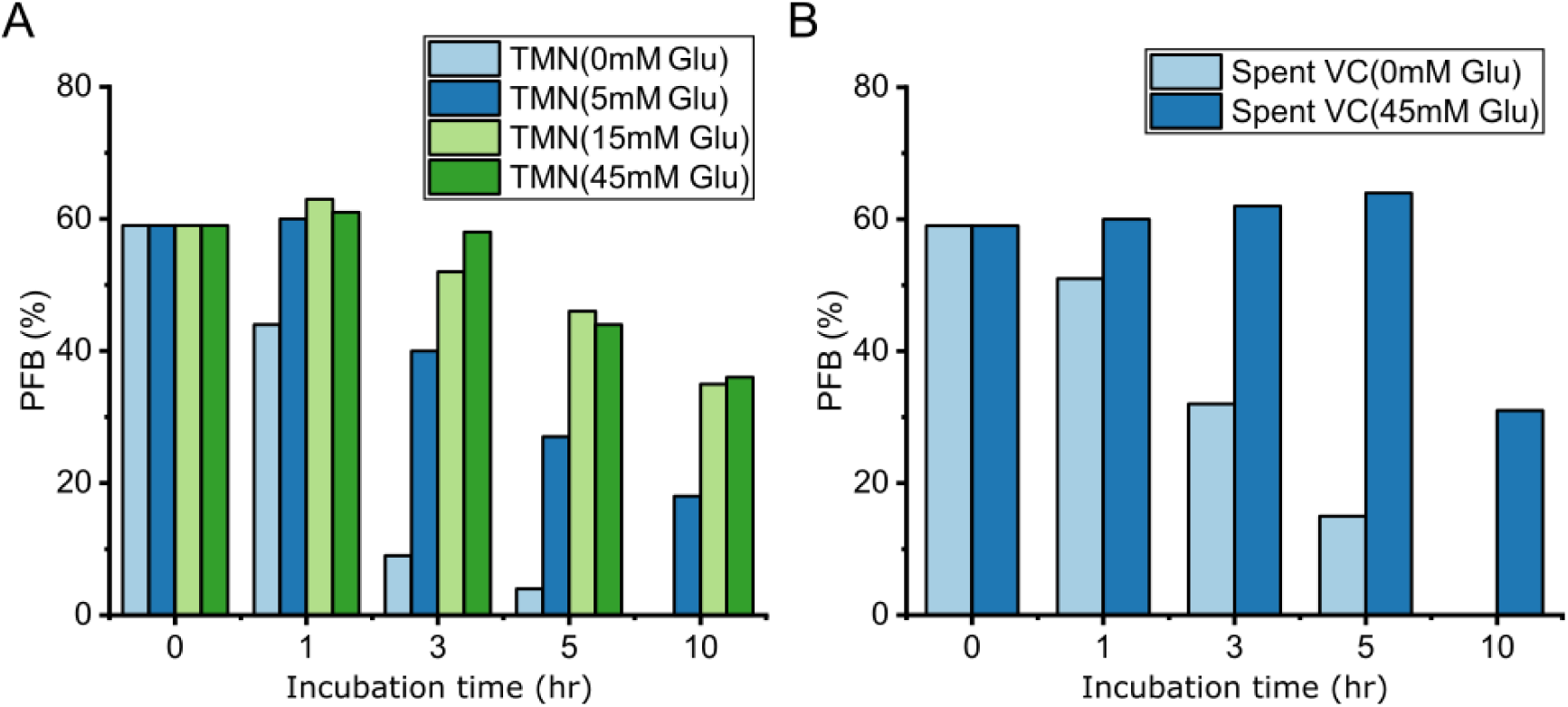
Lack of glucose promotes flagella loss. (A) Cells cultured for 3.5 hrs in VC medium were transferred to glucose-less TMN media supplemented with glucose in varying concentrations. (B) Cells cultured for 3.5 hrs in VC medium were transferred to spent cell supernatant with either no added glucose or supplemented with 45 mM glucose (At least 300 cells were counted for each time point.)

Besides the depletion of glucose, the accumulation of inhibitory metabolites in the stationary phase might also contribute to flagella loss. So, we prepared a ‘spent’ VC medium by removing bacterial cells from an overnight culture and used this supernatant to culture mid-exponential phase *V. alginolyticus* cells. As expected, the PFB showed an obvious decrease in the spent VC medium (Figure 4B); however, for cells transferred to the spent VC medium supplemented with 45 mM glucose, we found that the PFB would remain stable for 5 hrs before beginning a gradual decline. This indicated that glucose supplementation alone was sufficient to rescue flagellar loss in the spent VC medium. Taken together, our data demonstrated that the lack of a carbon source, glucose, is a primary factor driving active bacterial flagella abandonment.

### Fast movement of FliG clusters on the inner membrane before flagellar loss

To investigate the mechanics of flagellar loss in the stationary phase, we used two-color imaging to simultaneously record flagellar disassembly and FliG cluster movements. At first, we took time-lapse images every 20 mins to investigate whether the FliG cluster stayed with the bottom of the flagellum after flagellar disassembly. Interestingly, of the 72 cases of active flagella shedding we observed, in 69 cases (95.83%) the EGFP-FliG cluster was missing from the base of the flagellum before the flagellum detached from the cell body, while in the remaining 3 cases (4.17%) the EGFP-FliG cluster stayed in the cell bodies after the flagella detached (Figure 5A, B). Based on these results, we speculated 2 scenarios that may underlie the flagellar disassembly process: 1) The disassembly process starts with proteolysis of the basal body (C-ring), followed by MS-ring deconstruction, then flagellum detachment from the cell body. 2) The disassembly process starts from breakage somewhere in the rod connecting the flagellar filament with the basal body, causing both the flagellum and the C-ring to detach.

**Figure 5.**
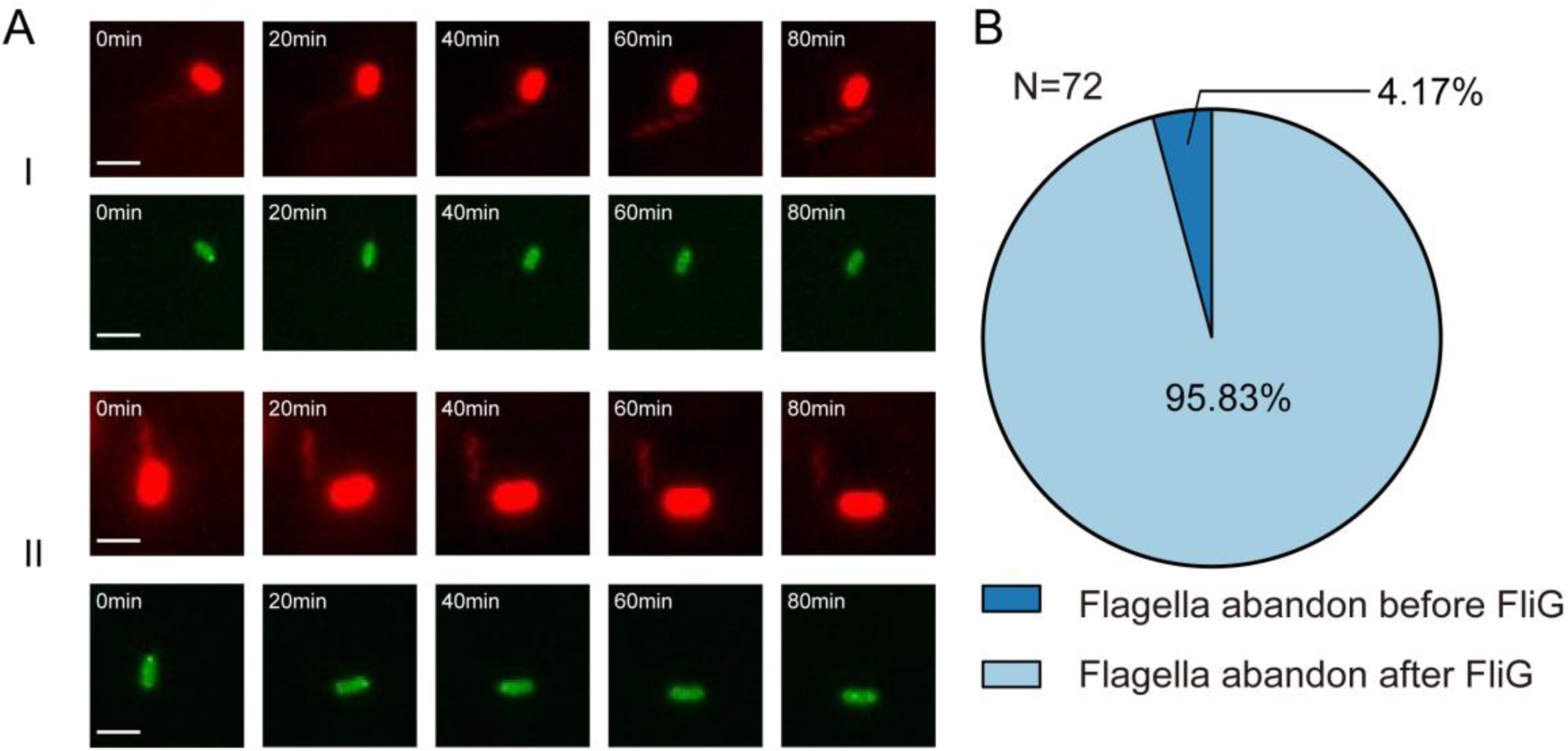
Two-color fluorescent images showing the flagellar disassembly process (A) Representative examples showing the FliG cluster was missing from the base when the flagellum detached from the cell body. I. II Two representative examples showing that flagellum was abandoned after FliG disappearance. (B) Percentage of FliG clusters missing from the base vs. not missing when the flagellum detached from the cell body

To differentiate these 2 possible mechanisms, we traced the movement of EGFP-FliG clusters with time-lapse imaging at 10 secs intervals during flagellar loss. Surprisingly, we saw that many polar, localized FliG clusters suddenly moved (Figure 6A). We described this movement using a custom computer program developed to track FliG cluster trajectory (see Material and Methods). Interestingly, moving FliG clusters were found to travel along the periphery, not the center, of bacterial cells (Figure 6B). Next, we measured the distances between mobile FliG clusters and the edges of bacterial cells. The distance distribution was found to be similar to that of localized FliG clusters to the edges of bacterial cells, suggesting that these mobile FliG clusters were moving on the cells’ inner membranes (Figure 6C). This finding also implied that flagellar loss starts with detachment, rather than destruction, of the C-ring complex. Next, we plotted the mean-squared displacement (MSD) versus time interval (Δt) of the mobile FliG clusters. The slope of the MSD-Δt plot confirmed that EGFP-FliG clusters were highly mobile with an average diffusion constant of 9.45×10^−3^ μm^2^/s (Figure 6D). In a previous study, Fukuoka et al. [44] showed that GFP–FliG clusters move on the inner membrane in *Escherichia coli* with an average diffusion constant of 4.9×10^−3^ μm^2^/s, thus supporting our premise that *V. alginolyticus* FliG clusters also move on the inner membrane before flagellar disassembly.

**Figure 6.**
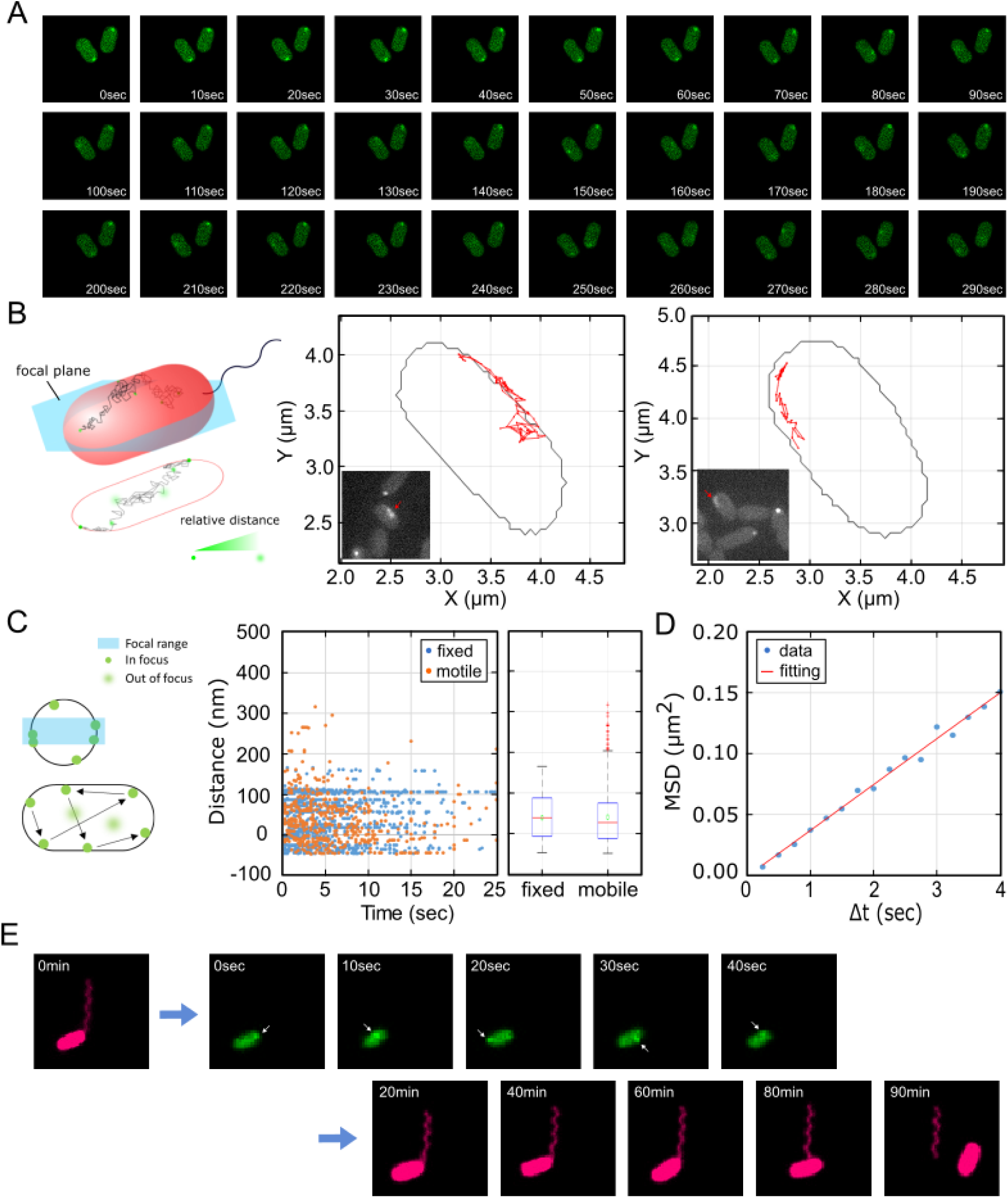
Fast diffusion of FliG clusters during flagella loss. (A) Time-lapse fluorescence images of representative *V. alginolyticus* (NMB344) EGFP-FliG in glucose-less TMN medium. Some FliG clusters that were localized at the cell poles suddenly moved away from the pole. (B) The focal plane is kept in the middle of cells during the tracking. Two example traces are shown here. The trajectory of a moving FliG demonstrated that FliG clusters traveled along the periphery of bacteria, implying that flagellar loss starts with FliG cluster removal, not FliG cluster proteolysis. None of observed FliG traces shows crossing event through cell center. Insets: Accumulated time-lapse fluorescence images corresponding to the graphics. (C) We analyzed the distance between FliG clusters and the cell edge as we keep focus on the cell middle plane. Although the motile FliG cluster is moving, it remains close to the cell edge as the distributions of distances between the edges is the same of localized FliG clusters. N _fixed_ = 22, N_motile_ = 35. (D) The linear fit of the MSD-Δt plot of FliG clusters revealed an ensemble average diffusion constant of 9.45×10^−3^ μm^2^/s. 27 cells were examined. (E) Dynamic fluorescence imaging of a bacterial flagellar filament (NMB344, red) and EGFP-FliG diffusion in glucose-less TMN medium. Arrows indicate location of FliG.

FliG is located on the cytoplasmic side of the C-ring, which is also directly associated with the MS-ring component, FliF [37]. The fast movements of FliG clusters that we observed suggested that the MS-ring and C-ring motor components can move freely on the inner membrane, after losing the anchoring forces of the rod, P-ring, and L-ring [45-47]. We further confirmed the connection between FliG movement and flagella loss using two-color time-lapse imaging (Figure 6E). The results clearly demonstrated that the FliG cluster definitely moves freely on the inner membrane before the flagellum detaches from the cell body. Altogether, our data suggest that flagellar loss is not initiated by basal body deconstruction but rather by breakage in the rod component somewhere between the flagellar filament and the C-ring (Figure 7).

**Figure 7.**
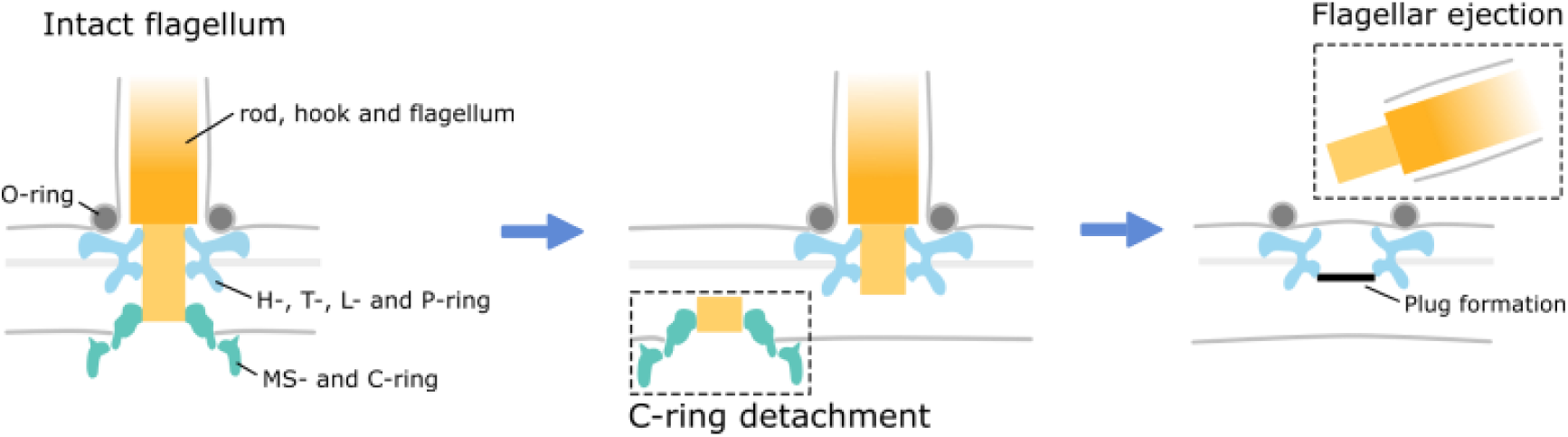
A model summarizing the disassembly process of a *V. alginolyticus* flagellum showing that it begins above the MS-ring before FliG depolymerization. The C-ring, with inner membrane components, then mobilizes on the cell membrane. Finally, the LP-ring is sealed and the flagellum ejected.

## Discussion

Using the single polar flagella of *V. alginolyticus* as our model, we found that the PFB in a population varies substantially across different growth phases and revealed that bacterial cells can stop making flagella, or even actively abandon flagella, upon nutrient depletion. To discover this, we used *in vivo* fluorescence imaging to systematically survey the dynamic production and loss of flagellar filaments, critical cellular structures for bacterial motility, during the bacterial life cycle.

Why bacterial cells choose to dynamically produce and lose flagellar filaments remains a mystery, especially since the production of flagellar filaments consumes substantial amounts of protein and energy. Instead, why don’t the bacteria produce the flagellar filaments, keep them as permanent cellular structures, and control when it should be used/powered? Several known mechanisms enable bacteria to stop flagellar rotation. For instance, under nutrient depletion, activation of cyclic di-GMP signaling triggers YcgR, a c-di-GMP binding protein, to interact with the flagellar switch-complex proteins FliG and FliM, stopping flagellar rotation and acting as a ‘molecular brake’ [48, 49]. In addition, decreasing the PMF has been shown to dissociate stator units from the flagellar motor, thus stopping motor rotation in *E. coli*. Similar observations have also been found in the sodium-driven motor in *V. alginolyticus* [50].

Given these existing mechanisms that enable bacteria to stop flagellar rotation without losing flagella, we were intrigued that they would actively abandon their flagella, as we observed during their prolonged stationary phase. Despite a comprehensive understanding of the flagellar assembly, the process of flagellar disassembly remains poorly understood. For *Caulobacter crescentus*, flagella ejection occurred during its growth cycle and previous studies showed that ejection started from the inside out [51], first with digestion of the MS protein, FliF. Recently, work using electron micrograph revealed that the *C. crescentus* disassembly product does not include the C-ring, MS-ring, and export apparatus [52]. However, Ferreira et al. [39] showed the ejected flagella in 5 γ-proteobacteria would leave relic structures composed of the P-, L-, H-, and T-rings and the basal disk in the outer membrane. All these findings came from biochemical and electron microscopic studies but lacked dynamic imaging of the disassembly process. In our work, we found that flagellar ejection in *V. alginolyticus* starts with a broken connection somewhere between the flagella and the MS-ring, leading to the fast movement of FliG clusters, which differs from the ‘inside out’ model of disassembly. Recently, Kaplan et al. [38] observed an inner-membrane sub-complex containing the C- and MS-rings near the P- and L-ring sub-complexes in many *Legionella pneumophila* cells. Although they explained this with a cell membrane breakage event, perhaps these disassembly products resulted from a ‘break in the rod’, as we observed and according to our proposed model. Furthermore, we established that flagellar loss is less affected by the newly produced substances in the stationary phase, but is more dependent on glucose in the medium, thus agreeing with a previous result [39]. In terms of pathogen-host interaction, whether the loss of flagella due to nutrient depletion would intensify or attenuate host immune response warrants continued study.

## Materials and Methods

### Strains and culturing

The *V. alginolyticus* wild-type strain VIO5 was used in this paper for flagellar visualization and the calculation of the PFB. The green fluorescent protein-FliG (EGFP-FliG) expressing strain (NMB344) was constructed from the VIO5 strain by using homologous recombination with the plasmid pTSK92, a derivative of the suicide vector pSW7848 [53], as described previously [54]. *V. alginolyticus* was grown from frozen stock in VC medium (0.5% Bacto-tryptone (w/v), 0.5% yeast extract (w/v), 0.4% K_2_HPO_4_ (w/v), 3% NaCl (w/v), 0.2% glucose (w/v)) for 16 hours. The cells were then re-grown in VC medium after 1:100 dilution. Two motility buffers, TMN (50 mM Tris-HCl (pH = 7.5), 5 mM MgCl_2_, 300 mM NaCl) and TMK (50 mM Tris-HCl (pH = 7.5), 5 mM MgCl_2_, 300 mM KCl), with selected glucose concentrations were used as non-growing media.

### Sample preparation

To visualize *V. alginolyticus* flagella, or FliG clusters, we constructed a micro-channel chamber by using double-sided tape to affix a coverslip to a microscope slide that had both been cleaned with saturated KOH solution. To immobilize cells, poly-L-lysine (Sigma, 0.1%) was run through the channel, which was then washed immediately with TMK medium. Then, the cells were harvested and washed with TMK by centrifuge with 5500 rpm for 2 mins. Next, we added the cell suspension into the channel, allowing it to sit for 10 mins before washing it with TMK to remove non-attached cells.

### Labeling flagella and measuring the PFB

At a selected time point, we labeled an aliquot (100 μL) of cells with 16 μM fluorescent dye FM 4-64 (T3166, ThermoFisher) in TMN. The labeled cells were then loaded into the micro-channel chamber and allowed to sit 30 min to ensure that all cells were immobilized on surfaces. To measure the PFB we used a Nikon Ti-E Eclipse inverted microscope equipped with a 1.45 NA 100x Plan Apo lambda oil objective, an sCMOS camera (Zyla 4.2 plus, Andor), an LED illumination light source (pE4000, CoolLED), a 530nm/30x bandpass center wavelength excitation filter, a 550nm longpass center wavelength emission filter, and the Nikon Perfect Focus system. Typically, we counted total 300-500 immobilized cells on both sides of the observation chamber of each time points.

For the glucose supplementation assay, cells were re-grown for 3.5 hrs then harvested by centrifuge with 5000 rpm for 10 mins. Then, the culture was incubated at 30 °C in TMN/Spent VC medium containing various glucose concentrations. Next, flagella are labeled and we counted the PFB using the above PFB measurement method.

### Time-lapse fluorescent microscopy

We observed time-lapse fluorescence imaging of flagella/FliG dynamics through an Olympus fluorescence microscope with a 1.46 NA 100x oil objective and using the Z drift compensation system at room temperature. FM4-64 was excited by a 561 nm laser (OBIS 561, Coherent) and observed through a 594 nm optical filter, while EGFP-FliG was excited by a 488 nm laser and observed through a 530 nm filter. The poly-L-lysine coating on the coverslips of the micro-channel chambers immobilized the cells and their flagella on the glass surface, enabling several hours of observation. We photographed time-lapse fluorescence images with an electron-multiplying charge-coupled device camera (EM-CCD, Evolve Delta, Photometrics) at 5 min or 10 min intervals depending on the bacteria’s growth rate.

### Analysis of FliG movement

We observed FliG dynamics on a Nikon Ti-E Eclipse inverted microscope which is equipped with a 1.45 NA 100x Plan Apo lambda oil objective, an EM-CCD camera (iXon Ultra 888-Life, Andor), an LED illumination light source (pE4000, CoolLED), a 470nm/20x bandpass center wavelength excitation filter, a 530nm/30x bandpass center wavelength emission filter, and the Perfect Focus system. To measure cell contours, we first photographed a phase-contrast image and then captured fast dynamic images of FliG clusters at 250 ms exposure time each for a 3 min duration using the Nikon NIS-Elements platform. Optical focus was kept in the middle plane of cells and cell contours were found using phase contrast images and the Otsu algorithm.

For FliG spots of interest, a custom-wrote MATLAB program was used to trace the spots’ trajectories. First, we subtracted the constant cell fluorescent signal to remove autofluorescence signals. Then, spots on the image were fitted by 2D Gaussian functions to define a center and a full width at half maximum. Finally, traces with non-focus spots were discarded.

### Detection of the filament protein from the culture supernatant

Overnight culture of the VIO5 cells was diluted 1:100 to the fresh VC medium, and cells were regrown at 30°C. At indicated time points, absorbance of 660 nm was measured and then 1 ml of the culture was aliquoted. Cells were precipitated by centrifugation (16,000 × *g*, 5 min), and the 900 µl of culture supernatant was ultracentrifuged (154,000 × *g*, 30 min). The pellet was suspended to the normalized volume (by OD_660_ of the culture) in the V buffer [50 mM Tris-HCl (pH 7.5), 300 mM NaCl and 5 mM MgCl_2_]. The pellet suspensions and whole cell lysates were analyzed by SDS-PAGE followed by the immunobloting as described previously [54]. The antipolar flagellin antibody (PF42) was used to detect flagellin [55].

## Acknowledgement

This work is financially supported by the Ministry of Science and Technology, Republic of China, under contract No. MOST-107-2112-M-008-025-MY3 to CJL and by the National Natural Science Foundation of China (No. 31722003, No.31770925) to FB, and by the Ministry of Education, Culture, Sport, Science and Technology of Japan (Grant number JP26115705) and Japan Society for the Promotion of Science (Grant number JP16H04774) to SK. FB and CJL are also supported by a Human Frontier Science Program grant (RGP0041/2015).

---

**Fig. S1.**
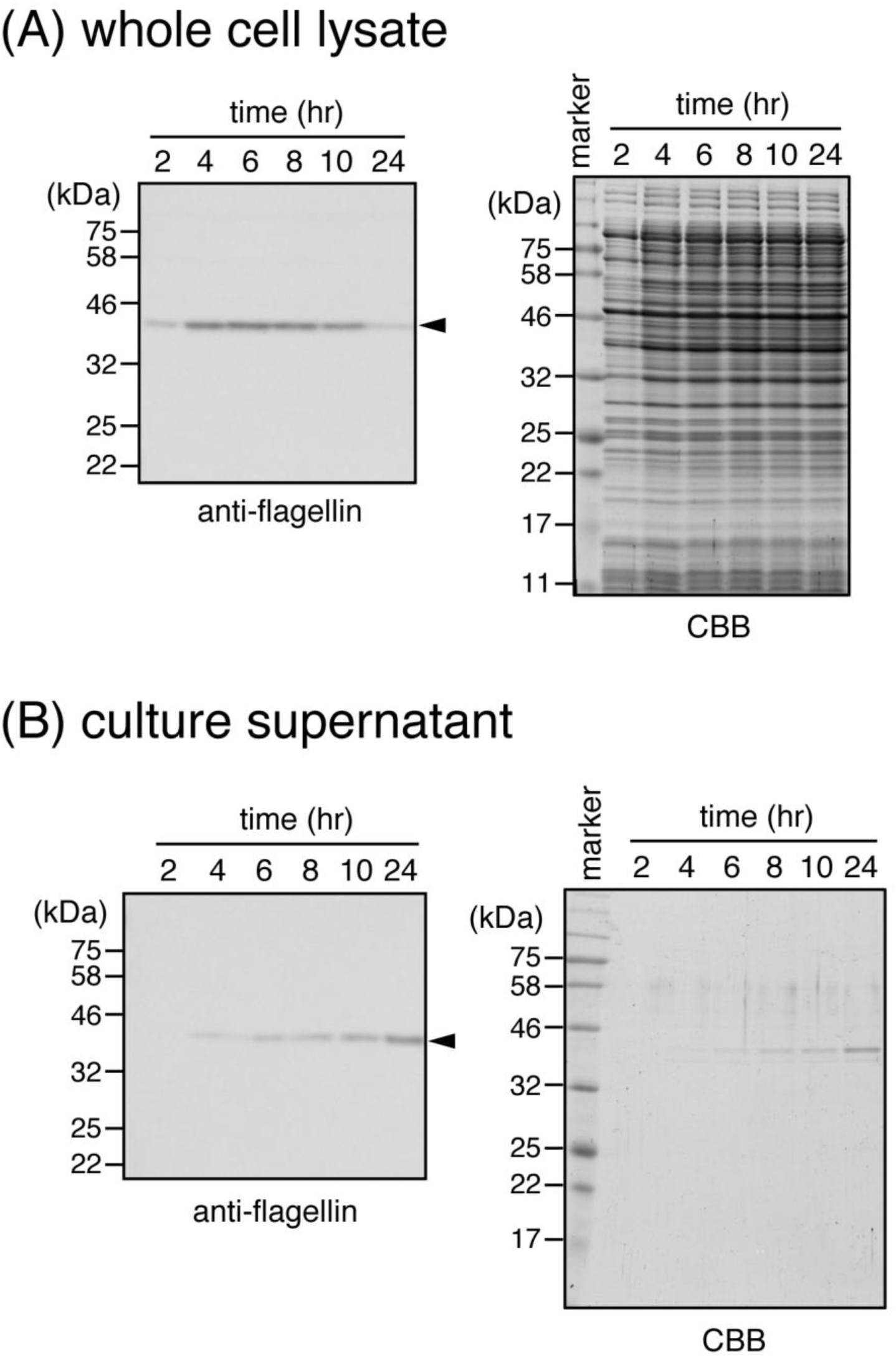
Detection of the filament protein in the culture supernatant. VIO5 cells were grown in VC medium and 1 ml culture was aliquoted at indicated time. Cells were precipitated by centrifugation, and then culture supernatant were ultracentrifuged. Both pellets (low speed spin for “whole cell lysate” and ultracentrifugation for “culture supernatant”) were suspended to the normalized volume (equivalent to OD_660_of 10) and analyzed by SDS-PAGE followed by immunoblotting. (A): analysis of whole cell lysate, (B): analysis of culture supernatant. Left panels show anti-flagellin immunoblot, and right panels show Coomassie Brilliant Blue staining of SDS-PAGE gel. Polar flagellin is indicated by the arrow head.

